# The sialylation pattern of dietary glycopeptides is vital for inhibiting pathogens and modulating gut microbiota in elderly individuals

**DOI:** 10.1101/2025.04.02.646947

**Authors:** Jiao Zhu, Tiantian Zhang, Jianrong Wu, Hongtao Zhang

**Author notes:** Corresponding author: *E-mail addresses:* (J. Wu).

## Abstract

N-glycosylation is a post-translational modification of proteins that can influence the structure and function of glycoproteins. Edible bird’s nest (EBN) contains highly sialylated mucins with N-glycans and O-glycans, while casein glycomacropeptide (CGMP) is modified with only O-glycans. We prepare EBN sialoglycopeptides through sequential protease treatment and their inhibitory effects on pathogens, including *Candida albican*s, *Helicobacter pylori*, and avian influenza virus (AIV), as well as their modulation of gut microbiota in elderly individuals, were compared with CGMP. N-glycan analysis revealed the presence of multiantenna N-glycans, characterized by sialylation and bisecting GlcNAc in the EBN glycopeptides. EBN glycopeptides significantly inhibited *C. albicans* and *H. pylori*. However, the removal of terminal sialic acid from EBN glycopeptides resulted in a marked reduction of this inhibitory effect. Using Maackia Amurensis Lectin II as an AIV mimic, EBN glycopeptides effectively inhibited AIV adhesion to MDCK cells. In vitro fecal fermentation using fecal samples from elderly individuals revealed that the EBN glycopeptide group exhibited remarkable probiotic effects on the gut microbiota, increasing the relative abundance of beneficial bacteria such as *Lactobacillus* and *Roseibacterium*, while decreasing the relative abundance of harmful bacteria such as *Escherichia-Shigella* and *Klebsiella.* Overall, EBN glycopeptide, rich in sialylated N-glycans and O-glycans, strongly inhibited pathogenic microorganisms and showed superior probiotic properties compared to CGMP with sole O-glycosylation.

## 1. Introduction

N-acetylneuraminic acid (Neu5Ac) is a negatively charged 9-carbon α-keturonic acid that plays crucial roles in neural development, immune regulation, pathogen adhesion and tumor metastasis, among other processes (Ling et al., 2022). Neu5Ac is the predominant form of sialic acid present in mammals. It is widely distributed in nature, with the highest concentrations occurring in the edible bird’s nest (EBN), casein glycomacropeptide (CGMP), and other biological sources. EBN refers to nests constructed by swiftlets, consisting of a mixture of secretions from their salivary glands (Chu et al., 2023). These EBNs have garnered significant attention due to their high nutritional value and market potential (Chok et al., 2021). Recent studies have demonstrated that EBN plays a critical role in alleviating neurodegenerative diseases (Yew et al., 2019), enhancing intestinal health (Wu et al., 2023), and providing whitening and anti-aging benefits (J. Wang et al., 2024). These diverse biological functions of EBN are attributed to its primary bioactive component, sialoglycoprotein, in which the bound Neu5Ac plays a key role (T. Zhang et al., 2023). Through enzymatic hydrolysis, the water-insoluble EBN sialoglycoproteins can be converted into water-soluble sialoglycopeptides (SGPs), which are readily applicable in cosmetics and beverages (T. Hui Yan et al., 2021; S. R. Ng et al., 2020). Casein glycomacropeptide (CGMP) is a bioactive peptide derived from milk, characterized by O-glycosylation and containing approximately 7% sialic acid. Numerous biological and functional properties of CGMP are attributed to the presence of terminal sialic acids (Y. Lu et al., 2022).

Glycosylation is a prominent post-translational modification of proteins, involving both N-glycans and O-glycans, which are indispensable for maintaining protein stability and supporting essential physiological functions (You et al., 2014). Over 50% of mammalian proteins undergo glycosylation, which is involved in a variety of important processes, including cell recognition and signaling, immune cell recognition, neuronal development and function, and regulation of cell growth (Pasala et al., 2024). Glycoproteins exhibit structural diversity due to variations in glycan composition, glycosylation sites, and the protein backbone (Čaval et al., 2021). N-glycosylation of proteins affects glycoprotein folding, transport, stability, molecular recognition, and interactions (L. Zhang et al., 2023). Many therapeutic proteins and enzymes undergo N-glycosylation, and their solubility, stability, safety, immunogenicity, and efficacy are directly related to the level of N-glycosylation and the structure of the N-glycans (Delobel, 2021). Dysregulation of N-glycosylation is associated with a range of diseases, including cancer, neurodegenerative diseases, infections, and inflammation. Furthermore, N-glycans can affect the infectivity of pathogen-associated proteins by modulating their conformation, thereby affecting their ability to infect human cells(Pasala et al., 2024). Several human dietary glycoproteins including ovomucin, EBN, egg IgY, are rich in sialic acid, primarily in the form of N-glycans or O-glycans (Unal et al., 2024). Nevertheless, the biological functions of N-glycosylation in dietary glycoproteins remain to be fully elucidated.

Sialoglycans can inhibit pathogens, including *Candida albicans*, *Helicobacter pylori*, and avian influenza virus (AIV) by binding to their surface receptors. *C. albicans* is a significant opportunistic pathogen commonly found in the human gastrointestinal tract and oral cavity (Ramage et al., 2005). Mucins, which are heavily sialylated glycoproteins, downregulate genes involved in *C. albicans* invasion, thus preventing its attachment to host cells and the formation of biofilms (Kavanaugh et al., 2014). Moreover, the glycan chains present at the mucin O-glycosylation site can bind to surface proteins of *C. albicans*, thereby interfering with their attachment and invasion into host cells (Takagi et al., 2022). The glycoprotein derivative lactoferrin-derived LfcinB15 (N-glycosylation with or without sialic acid), exhibits potent bactericidal activity against *C. albicans* by disrupting its cell membrane and wall, thus inhibiting its growth and reproduction (C. K. Chang et al., 2021).

*H. pylori* is a prevalent bacterial pathogen responsible for significant gastrointestinal morbidity globally, and it is associated with peptic ulcers, gastritis, and MALT lymphoma (C. S. Chang et al., 2012). *H. pylori* can bind to and colonize mucins on the surface of the oral and gastric mucosa, with sialic acid-binding adhesin (SabA) playing a crucial role (Linden et al., 2008). A study by Yang et al. demonstrated that the combination of catechins and sialic acid could eliminate *H. pylori* infection by attenuating the Caspase-1 signaling pathway in gastric epithelial cells (J. C. Yang et al., 2013). Additionally, lactoferrin can also inhibit the secretion of *H. pylori* under iron-limited conditions, thereby exerting antibacterial effects (J. Lu et al., 2021). It is well-known that sialic acid plays a crucial role in the invasion of AIVs (Spackman, 2014). AIV enters animal cells and causes infection by binding hemagglutinin (HA) on the virus surface to sialic acid on the mucin terminals of host cells, which is subsequently released by viral neuraminidase (Stencel-Baerenwald et al., 2014; X. Yang et al., 2014). Current therapeutics for AIV infection, such as oseltamivir and zanamivir, are structural analogs of sialic acid, which exert their effects by inhibiting neuraminidase. Recent studies have shown that dietary sialoglycans, such as EBN and ovomucin, exhibit promising inhibitory effects against AIV. This is primarily attributed to their sialic acid content, derived from sialylated glycoproteins or glycopeptides that bind to hemagglutinin on the virus surface (Haghani et al., 2016; Haghani et al., 2017; Q. Xu et al., 2018). Furthermore, human intestinal microorganisms plays an important role in maintaining human health, and the presence of sialic acid in the gastrointestinal mucosa influences the composition of gut microbiota, pathogen colonization, and the intestinal barrier (Bell et al., 2023). Compared with young individuals, the abundance of intestinal flora (including probiotics such as *Bifidobacteria*) in the elderly is reduced, resulting in increased susceptibility to metabolic disorders and chronic inflammation (H. J. Yu et al., 2020). Effective interventions on gut microbiota may be important for delaying aging and preventing disease in the elderly. Previous studies have proved that EBN can increases the abundance of probiotics in the gut microbiota of pregnant women (Wu et al., 2023).

This study compared the inhibitory effects of EBN glycoproteins and CGMP on the growth of *C. albicans*, *H. pylori*, and AIV, and investigated their capacity to modulate the intestinal flora in elderly individuals. As controls, desialylated EBN sialoglycoprotein (Desia-E), sialic acid monomer (Neu5Ac) and 3’-sialyllactose (3’-SL) were employed. Furthermore, the N-glycan structures of the EBN SGP were profiled to better elucidate the associations between its structure and function.

## 2. Materials and methods

### 2.1 Materials and strain

EBN was supplied by Xiamen Yan Palace Seelong Biotechnology (Xiamen, China). The Bradford protein assay kit was obtained from Beyotime Biotechnology (Shanghai, China). Pronase was acquired from Yuanye Biotechnology (Shanghai, China). Trypsin was obtained from Scigrace Biotechnology (Shanghai, China). Alkaline protease was acquired from Solarbio Life Science (Beijing, China). Maackia Amurensis Lectin II (MAL-II) was acquired from Vector Labs (Burlingame, CA, USA). Sulfonated cyanine 5-N-hydroxysuccinimide ester was obtained from Duofluor, Inc. (Wuhan, China). 3’-Sialyllactose (3’-SL) was generously donated by Chr. Hansen (Denmark), while casein glycomacropeptide (CGMP) was supplied by Arla Food Ingredients (Denmark).

*C. albicans* (ATCC 10231) and *H. pylori* (ATCC 43504) were provided by Guangdong Microbial Culture Collection Centre (Guangzhou, China). MDCK cells were obtained from Wuhan Pricella Biotechnology (Wuhan, China), while minimum essential medium (MEM) (HyClone) was obtained from Cytiva, USA, and fetal bovine serum was obtained from Gibco, USA. The brain heart infusion agent and Columbia blood agar were obtained from Thermo Fisher Scientific. Yeast malt (YM) agar medium, liquid Sartorius medium, *H. pylori* additive, and Brinell’s broth were purchased from Qingdao Haibo Biotechnology (Qingdao, China). The composition of basal nutrient culture media for in vitro fermentation of fecal bacteria was adopted from the reference (D. Yu et al., 2023). The remaining reagents were purchased from Sinopharm Chemical Reagent Co. (Shanghai, China).

### 2.2 Preparation of EBN glycopeptides

EBN glycopeptides were prepared through cascaded enzymatic hydrolysis, as illustrated in Fig. 1. The EBN pretreatment method was based on the work of Guo et al. with modifications (Guo et al., 2006). The dried EBN was rinsed three times with deionized water, followed by soaking in deionized water (5 g of EBN/300 mL) and left overnight at 4°C for 16 h. The samples were subsequently boiled for 30 min, freeze-dried to yield lyophilized EBN powder, and stored in a dry dish at room temperature.

**Fig. 1.**
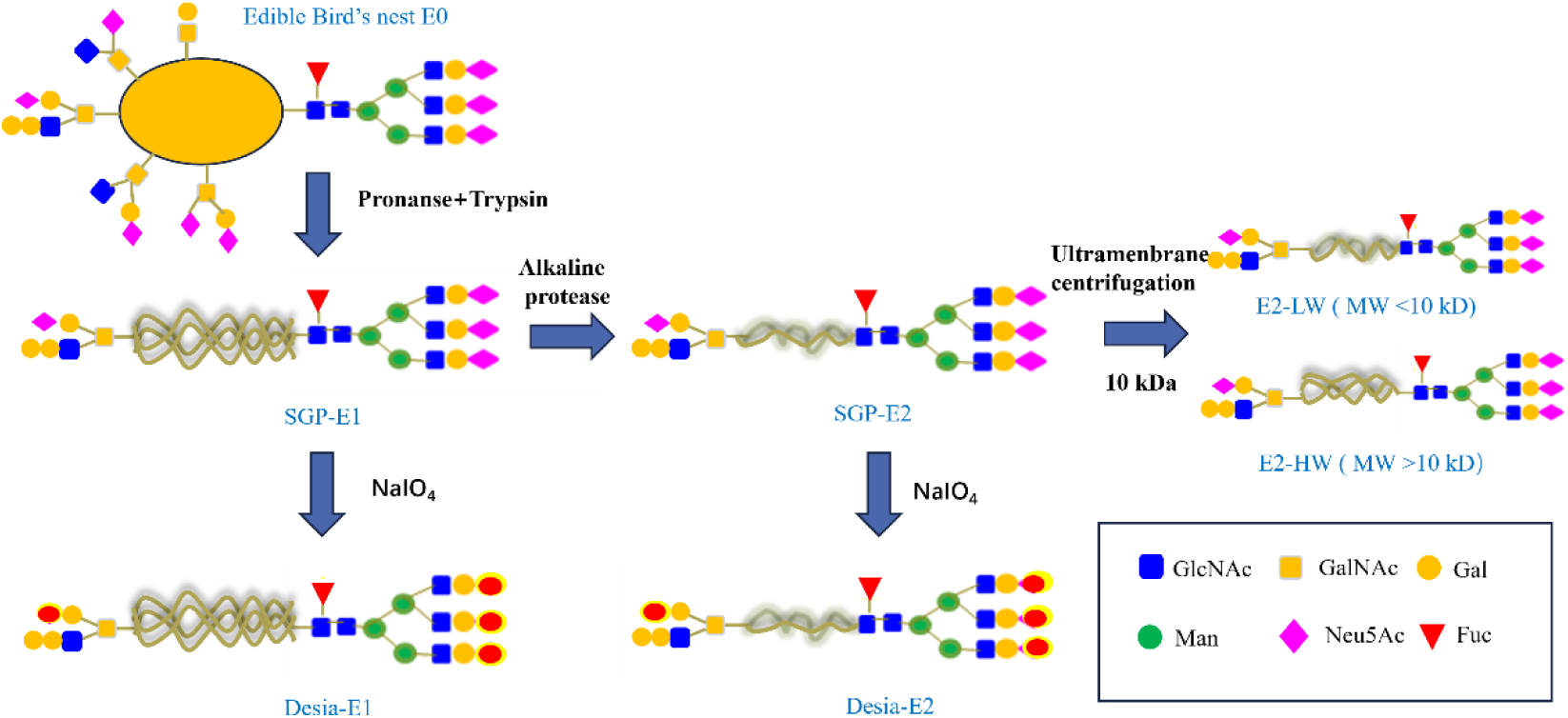
Preparation of edible bird’s nest sialoglycopeptide

The hydrolysis of EBN using proteinase was conducted in two stages. The objective of the first stage was to maximize the solubility of the protein. The protocol for this stage was adapted from the literature, with some modifications that included both double enzyme digestion and single enzyme digestion (Ghassem et al., 2017; Khushairay et al., 2014; Oda et al., 1998). The resulting digested sample was labeled as SGP-E1. The enzymatic combinations for the two-enzyme digestion groups included pepsin + trypsin, pancreatin + alkaline protease, and pronase + trypsin.

For the pepsin + trypsin treatment, EBN (10 mg/mL) was mixed with potassium chloride-hydrochloric acid buffer (0.1 M, pH 2.0). Pepsin (1%, w/w) was added to the mixture, which was then incubated at 37°C with continuous shaking for 2 h. Subsequently, the pH of the mixture was adjusted to 8.0 with 1 M NaOH, followed by the addition of trypsin (1%, w/w). The mixture was further incubated at 37°C for an additional 2 h. The enzymatic reaction was terminated by heating the sample in a boiling water bath for 15 min. After cooling to room temperature, the EBN hydrolysate was centrifuged at 4°C and 8000 rpm for 15 min. The supernatant was collected and subsequently freeze-dried. For the pancreatin + alkaline protease treatment, EBN (10 mg/mL) was mixed with potassium phosphate buffer (0.1 M, pH 8.0). Pancreatin (1%, w/w) and alkaline protease (1%, w/w) were subsequently added to the mixture. The reaction mixture was shaken for 2 h at the respective temperatures (pancreatin: 38°C, alkaline protease: 60°C). For the pronase + trypsin treatment, pronase (1%, w/w) and trypsin (1%, w/w) were used for hydrolysis for 2 h each. The reaction conditions for pronase (39°C, pH 8.0) and trypsin (39°C, pH 8.0) were maintained throughout the enzymatic treatment.

The second stage aimed to obtain EBN glycopeptides with a lower molecular weight (C. H. Ng et al., 2022). Papain (1%, w/w), alkaline protease, pancreatin, and pepsin were used to treat SGP-E1, resulting in the formation of SGP-E2. The optimal conditions for the papain reaction were a pH of 6.0, a temperature of 60°C, and a reaction time of 2 h.

### 2.3 Preparation of desialylated EBN glycopeptides

The EBN glycopeptide was reacted with sodium periodate to yield a desialylated glycopeptide (D. S. Kim et al., 2014). A total of 900 μL of SGP-E1 or SGP-E2 (10 mg/mL) was mixed with 100 μL of a NaIO_4_ solution (100 mM). The mixture was then incubated at 4°C for 30 min. After the reaction, the mixture was transferred to a 1000 Da dialysis bag and dialyzed for 3 days to remove any unreacted NaIO_4_. The dialysate was collected and subsequently freeze-dried to obtain desialylated samples (Desia-E1, Desia-E2).

### 2.4 Determination of sialic acid

The sialic acid content in EBN glycoproteins was determined using high-performance liquid chromatography (HPLC) (Martin et al., 2007; You et al., 2014). Initially, sialic acid was released from EBN glycoproteins by treatment with 4 mol/L acetic acid in a water bath at 80°C for 3 h. The sialic acid was then derivatized with o-phenylenediamine (OPD). The OPD derivatives were prepared by dissolving 0.01 g of OPD in 1 mL of 0.2 M sodium bisulfite solution, followed by thorough mixing. Sialic acid content was quantified using an Agilent 1260 series HPLC (FLD detector) with a unitary C18 column (4.6×250 mm, 5 μm), an FLD excitation wavelength of 373 nm, and an emission wavelength of 448 nm (Agilent, USA). The mobile phase consisted of ultrapure water, acetonitrile, and methanol in a ratio of 85:8:7 (v/v/v) at a flow rate of 0.9 mL/min.

### 2.5 Physicochemical characterization of the EBN glycoprotein

#### 2.5.1 Soluble protein

The soluble protein content in the EBN hydrolysate samples was determined using a Bradford protein assay kit (Beyotime Biotechnology, Shanghai, China)(Tan Hui Yan et al., 2022). Bovine serum albumin was used as the standard in a 96-well plate, and 5 μL of protein sample was added to each well. Subsequently, 250 μL of G250 staining solution was added, and the mixture was incubated for 5 min. The absorbance at 595 nm was measured using a microplate reader (Synergy H4, Biotec, USA).

#### 2.5.2 Total sugar

The total sugar content in the EBN hydrolysate was determined using the phenol‒sulfuric acid method(Fang et al., 2024). In brief, 1.0 mL of the sample (10 mg/mL) was mixed with 1.0 mL of 5% phenol solution. Then, 5.0 mL of concentrated sulfuric acid was added, and the mixture was incubated for 5 min at room temperature. Subsequently, 200

μL of the reaction mixture was transferred to a 96-well plate, and the absorbance at 490 nm was measured using a microplate reader (Synergy H4, Biotec, USA).

#### 2.5.3 EBN hydrolysis rate

The enzymatic hydrolysis rate of the EBN glycoproteins was calculated via formula (1), which represents the ratio of the mass of the dried EBN sample after the first hydrolysis stage to the mass of the dried EBN sample prior to proteinase treatment.

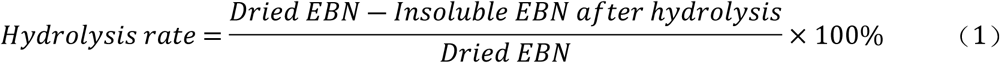

#### 2.5.4 Molecular weight

The molecular weight of the EBN hydrolysates were determined using high-performance gel permeation chromatography (HPGPC) (K. Wang et al., 2021). EBN SGPs were dissolved in ultrapure water (5 mg/mL) and subsequently filtered through a 0.22 μm filter membrane. HPGPC was conducted using a Wyatt DAWN HELEOS 8+ system (Waters, USA) equipped with a TSKGgel G3000SWXL column (7.8×300 mm) (Merck KGaA, Germany). The mobile phase consisted of 0.1 M sodium sulfate and 0.1 M sodium phosphate buffer solution (pH 6.7), with a flow rate of 0.5 mL/min.

### 2.6 Inhibitory effects of EBN glycopeptides on C. albicans

*C. albicans* was cultured on YM agar plates and in liquid Sartorius medium at 37°C. The culture mixture was adjusted to 3×10^8^ CFU/mL using a McFarland turbidity standard, followed by dilution to 1×10^5^ CFU/mL.

The inhibition assay was conducted following the method described by Kim et al. with slight modifications (Y. G. Kim et al., 2022). SGP-E2 was diluted in distilled water to final concentrations of 1 mg/mL, 2 mg/mL, 10 mg/mL, and 20 mg/mL. The controls included SGP-E1, Desia-E1, Desia-E2, E2-LW, and E2-HW, as well as sialic acid monomers (Neu5Ac), 3’-SL, and CGMP, each at a concentration of 20 mg/mL.

To prevent moisture dissipation from the inner wells during incubation, 200 μL of sterile water was added to outermost wells of a sterile 96-well plate. Subsequently, 100 μL of the prepared bacterial suspension was added to the inner wells, followed by 100 μL of SGP-E2 to achieve final concentrations of 0.5 mg/mL, 1 mg/mL, 5 mg/mL, and 10 mg/mL. Additionally, 100 μL of E1, Desia-E1, Desia-E2, E2-LW, E2-HW, Neu5Ac, 3’-SL, or CGMP (20 mg/mL) was added to the plate. Blank controls were prepared by adding 100 μL of sterile water to the remaining wells. Each sample was tested in triplicate. The 96-well plate was incubated in a shaker at 37°C and 120 rpm for 24 h, after which the OD_600_ value was measured.

### 2.7 Inhibitory effects of EBN glycopeptides on H. pylori

#### 2.7.1 Medium and culture conditions

The brain heart infusion medium was prepared by dissolving 9.6 g of brain heart infusion powder and 23.4 g of Columbia agar in 780 mL of distilled water, followed by autoclaving. Once the temperature reached 46°C, aseptic defibrillated sheep blood (33 ∼ 40 mL) and 1% (v/v) *H. pylori* additives were added.

For the Brinell broth medium, the pH was adjusted to 7.0 according to the instructions, and the mixture was then autoclaved at 121°C for 15 min. After cooling to room temperature, 5% (v/v) fetal bovine serum and 1% (v/v) *H. pylori* additives were added.

Cultivation was performed by plating 100 μL of *H. pylori* bacterial broth onto blood agar plates with brain heart infusion medium, followed by incubation at 37°C under microaerobic conditions (85% N_2_, 10% CO_2_, 5% O_2_) for 5 to 7 days. The activated colonies were used to inoculate 50 mL of Brinell’s broth medium, which was then incubated at 37°C with gentle aeration at 160 rpm for 3 to 5 days. The cell suspension was then centrifuged at 5000 rpm for 5 min, after which the cells were resuspended in 0.8% (w/v) saline. The suspension was adjusted to a density of 1×10^6^ CFU/mL using a McFarland turbidity standard and Brinell’s broth medium.

#### 2.7.2 Inhibition of H. pylori

The method used to evaluate the inhibitory effect on *H. pylori* was adapted from Wang et al. (J. Wang et al., 2020). Lyophilized EBN SGP (20 mg/mL) was added to the broth medium, and SGP-E1 and SGP-E2 were diluted to final concentrations of 1 mg/mL, 2 mg/mL, and 10 mg/mL, respectively. Additionally, Neu5Ac, 3’-SL, and CGMP (20 mg/mL) were independently added to Brinell’s broth medium.

A sterilized 96-well plate was filled with 100 μL of the prepared *H. pylori* bacterial suspension. Then, 100 μL of Brinell’s broth medium with or without the sialoglycan samples was added. Three replicates were prepared for each sample. The 96-well plates were incubated at 37°C in a microaerobic environment with shaking at 160 rpm for 72 h. At the conclusion of the incubation period, the OD_600_ value of each well was measured, and the inhibition rate (%) was calculated via formula (2).

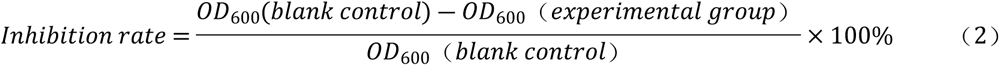

### 2.8 Inhibition of avian influenza virus by EBN glycoproteins

#### 2.8.1 Fluorescent labeling of the lectin MAL-II

A solution containing 1 mg of Cyanine5-N-hydroxysuccinimide (Cy5-NHS) in 1.0 mL of DMSO was incubated for 1 hour at room temperature in the dark. In a 5 mL centrifuge tube, 0.5 mL of Maackia Amurensis Lectin II (MAL-II) (2 mg/mL) was mixed with 0.06 mL of activated Cy5-NHS fluorescent dye and allowed to react for 3 h at room temperature. Following the reaction, the unreacted fluorescent dye was removed through ultrafiltration centrifugation using a 2 mL (10 kDa) ultrafiltration centrifuge tube and a 10 mM HEPES buffer solution. The purified Cy5-labeled MAL-II was stored at 4°C.

#### 2.8.2 Inhibition of MAL-II binding to MDCK cells by EBN glycoproteins

The ability of the EBN glycoproteins to inhibit the binding of AIV mimics to MDCK cells was assessed according to the method described by Wang et al. (X. Wang et al., 2021). MDCK cells were cultured in complete MEM at 37°C in a CO_2_ incubator with 5% CO_2_. When the MDCK cells reached 80% confluence, they were collected, centrifuged, resuspended, and counted. The cell density was adjusted to 10^4^ cells/mL, and the cells were seeded into confocal culture dishes. The cells were then incubated at 37°C in a CO_2_ incubator with 5% CO_2_ until they reached 60% ∼ 70% confluence. The cells were washed with PBS, fixed with 1 mL of 4% paraformaldehyde, and blocked with PBS containing 5% BSA for 1 hour at room temperature. Cy5-labeled MAL-II and various sialoglycan samples were added to the cell cultures and incubated overnight at 4°C in the dark. Subsequently, the MDCK cells were stained with DAPI staining solution, and images were captured using a laser confocal microscope (LSM710, Carl Zeiss AG, Germany) with the DAPI and Cy5 channels. Fluorescence intensity was analyzed using ImageJ software (National Institutes of Health, USA).

### 2.9 In vitro fermentation with intestinal microorganism

#### 2.9.1 Simulated digestion in vitro

The in vitro simulated digestion of the samples was conducted according to the protocol outlined in reference with slight modifications (Brodkorb et al., 2019). The in vitro simulated digestion consists of three stages, simulated salivary digestion, gastric digestion and intestinal fluid digestion.

In the simulated salivary digestion phase, 1.0 g each of E1, Desia-E1 and CGMP lyophilized powders was mixed with 25 mL of artificial saliva. The mixture was then incubated in a water bath at 37°C for 15 min with continuous shaking at 100 rpm. Artificial saliva without sample served as a blank control. The simulated salivary digestion process was maintained for 15 min, after which the enzyme was inactivated by boiling in a water bath for 10 min to terminate the digestion process.

For the simulated gastric fluid digestion phase, the pH of the simulated salivary digestive solution was adjusted to approximately 2.0 using 0.1 M HCl. An equal volume of artificial gastric fluid was then added, and the mixture was incubated in a water bath at 37°C for 120 min with continuous shaking at 100 rpm. The simulated gastric fluid digestion was conducted for 120 min, followed by enzyme inactivation through boiling in a water bath for 10 min to terminate the digestion process.

For the simulated intestinal fluid digestion phase, the pH of the simulated gastric fluid digestive solution was adjusted to about 7.0 using 1 M sodium bicarbonate. An equal volume of artificial intestinal fluid was then added, and the mixture was incubated in a water bath at 37℃ for 180 min with continuous shaking at 100 rpm. The simulated intestinal fluid digestion was carried out for 180 min, followed by enzyme inactivation through boiling in a water bath for 10 min to terminate the digestion process. The final digestive fluid obtained was the end product of the digestion process.

The Neu5Ac and 3’-SL were directly dissolved in the blank digestion solution at a concentration of 10 g/L, as they are not susceptible to digestion by proteinase or amylase.

#### 2.9.2 Volunteer feces collection and disposal

Volunteers were required to meet specific eligibility criteria for fecal donation, including the absence of biologic interventions, such as prebiotics and antibiotics, within the past two months, no history of gastrointestinal or chronic underlying diseases, and an age above 60 years. Fecal samples were collected from six healthy elderly donors using sterile fecal collection tubes. The samples were immediately stored in an ice box containing an anaerobic gas-producing bag, and processed within 6 h using the homogenization method (Aguirre et al., 2015). The fecal samples from the six donors were combined in equal proportions, diluted with sterile saline to a 10% volume fraction, and filtered through sterile gauze to remove food residues to obtain a fecal homogenate.

#### 2.9.3 In vitro anaerobic fermentation

Fecal bacteria were cultivated in vitro as described in reference (J. Xu et al., 2021). EBN digest samples, including E1, Desia-E1 and CGMP, were added to the basal medium at a final concentration of 5 g/L. Neu5Ac and 3’-SL digests were similarly added to the basal medium at a final concentration of 5 g/L. Blank digests were added to the basal nutrient medium as a control, with the unfermented fecal sample designated as the initial condition. Triplicate experiments were conducted for each group, and the fecal homogenate was inoculated at a 10% (v/v) concentration. In vitro fermentation was carried out in an anaerobic incubator at 37°C for 24 h.

#### 2.9.4 Organic acids analysis

One mL of fecal fermentation supernatant was sampled and 10 μL of internal standard (100 mM 2-methylbutyric acid), 250 μL of HCl, and 1 mL of anhydrous ether were added. The mixture was well-mixed, left at 4℃ overnight, and the transparent organic phase of the upper layer was collected. The sample was then dehydrated with anhydrous sodium sulfate and filtered through a 0.22 μm filter membrane. A gas chromatography system (7890A, Agilent, USA) equipped with an HP-INNOWas column (Agilent, USA) was used to determine the content of short-chain fatty acids (SCFAs) in the filtrate. The chromatographic conditions were as follows: the injector temperature was 220℃, the detector temperature was 250℃, the injection volume was 5 μL, the flow rate was 1.5 mL/min, the carrier gas was nitrogen, and the split ratio was 1:20. The contents of acetic acid, butyric acid and isovaleric acid were quantified using the internal standard method. The lactic acid content in the supernatant of fermentation broth was determined using a biosensor (SAB-40E, Shandong, China).

#### 2.9.5 Amplification analysis of intestinal microbiota

Genomic DNA was extracted from the fermentation broth using HiPure Stool DNA Kits (D3141, Guangzhou Meiji Biotechnology Co., China), and the V3 and V4 regions of the 16S rRNA gene were amplified with specific primers 341F and 806R, which included barcodes. Sequencing libraries were constructed using the Illumina DNA Prep Kit (Illumina, CA, USA). Library quality was assessed using the ABI StepOnePlus Real-Time PCR System (Life Technologies, USA), and sequencing was performed on the Novaseq 6000 (NovaSeq6000 S2 Reagent Kit v1.5, Illumina, USA) using the PE250 mode pooling technique. The sequencing results were filtered, corrected and assembled using DADA2 software (Illumina, USA) to obtain the valid data, including ASV (Amplicon Sequence Variant) sequence and ASV abundance information, which were subsequently analyzed.

### 2.10 Profiling of the N-glycans of EBN SGP

N-glycosylation is a critical post-translational modification of proteins that plays a significant role in the function of glycoproteins during host‒pathogen interactions and immune responses (Nagels et al., 2011). DNA sequencer-assisted fluorophore-assisted carbohydrate electrophoresis (DSA-FACE) is a highly sensitive glycomics detection method, enabling noninvasive and rapid high-throughput detection and analysis of N-glycan structures (Laroy et al., 2006). In this study, the N-glycans of EBN SGP were released through PNGase F treatment and labeled with 8-amino-1,3,6-pyrenetrisulfonic acid (APTS). N-glycan profiling was performed using a Glycome316 DSA-FACE GlycoDecoder (Superyears, Nanjing, China). The N-glycan database of plasma glycoproteins (Superyears) was set as the standard every run.

### 2.11 Statistical analysis

Each sample was analyzed in triplicated, and data analysis was conducted using Origin (2021) and GraphPad Prism 8 software. Significance was assessed using the Waller‒Duncan test with IBM SPSS Statistics 26 software. One-way analysis of variance (ANOVA) was used, and differences were considered statistically significant at *P < 0.05*, denoted by letters such as a, b, etc. The experimental data are presented as means ± standard deviations.

## 3. Results

### 3.1 Preparation and characterization of EBN glycopeptides

The hydrolysis of EBN by protease treatment presents significant challenges due to the extensive cross-linking of cysteine residues through disulfide bonds (Najafian & Babji, 2012). In our preliminary experiments, we found that EBN glycopeptides can be obtained through a two-stage protease treatment. As shown in Fig. 2A, the degradation of EBN by proteases varied significantly markedly when treated individually. Treatment with pepsin, trypsin, and pancreatin resulted in increased solubility of the protein, with the highest solubility observed when EBN was treated with a combination of pronase and trypsin. Therefore, the combination of pronase and trypsin was selected for the first stage of EBN hydrolysis, and the resulting hydrolysate was lyophilized and designated as SGP-E1. The first-stage hydrolysis yield was approximately 90.25%. Table 1 presents the sialic acid content of SGP-E1 as 11.57%, with the total sugar content measured at 16.57%. Notably, the chemical composition of EBN may vary depending on its origin and species. Our findings are consistent with the work of Daud et al., who reported that the protein fraction of EBN ranges from 59.8% to 66.9%, with the sialic acid content approximately 10% (Daud et al., 2021). The molecular weight distribution of soluble glycoproteins in EBN SGP-E1 was determined using HPGPC, as illustrated in Fig. 2C and 2D. The majority of SGP-E1 molecules exhibit a molecular weight greater than 10 kDa. In subsequent experiments, SGP-E1 underwent second-stage protease digestion, as depicted in Fig. 2C, and the hydrolysate was lyophilized and designated as SGP-E2. During alkaline protease digestion, SGP-E2 presented a relatively high concentration of glycopeptides with low molecular weights. Approximately 64.25% of glycopeptides in the alkaline protease digestion group (Fig. 2D) weighed less than 10 kDa, followed by the pepsin digestion group (59.14%), the pancreatic enzyme digestion group (40.23%), and the papain digestion group (21.52%). Consequently, alkaline protease was selected for the second-stage digestion of SGP-E1 to obtain the EBN glycopeptide, which was subsequently designated as SGP-E2. SGP-E2 was separated into a low-molecular-weight portion (E2-LW) and a high-molecular-weight portion (E2-HW) using a 10 kDa ultrafiltration centrifuge tube. To destroy the sialic acid in SGP-E1 and SGP-E2, sodium periodate treatment was applied. As illustrated in Fig. 2B, the desialylated sample, Desia-E1, exhibited a sialic acid content of 0.0917%, whereas SGP-E1 contained 11.57% sialic acid.

**Fig. 2.**
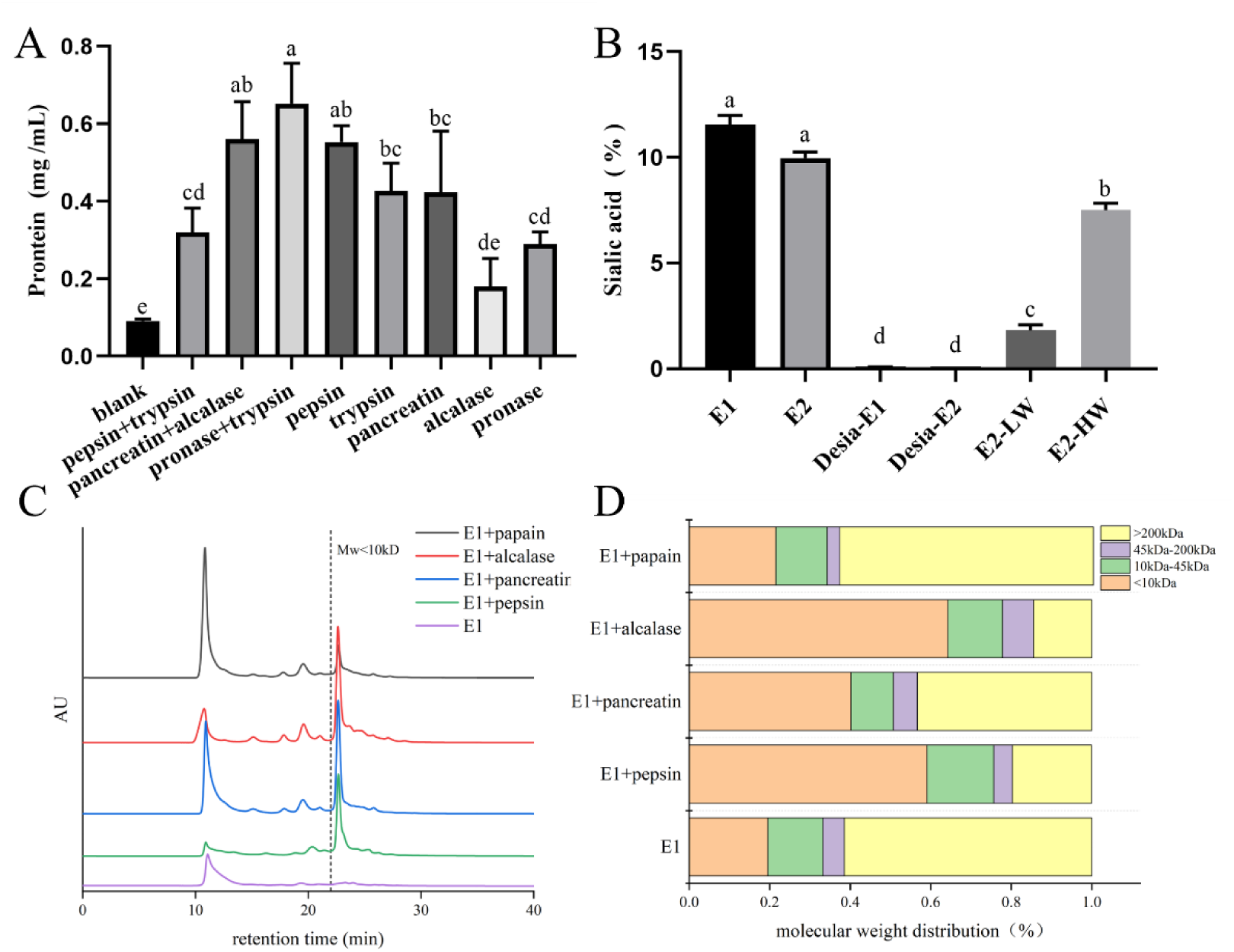
Effects of different proteases on EBN hydrolysis. Note: (A) Soluble protein content of EBN SGP-E1; (B) Sialic acid content of the EBN SGP; (C) EBN SGP-E2 preparation and molecular weight distribution; (D) Molecular weight of EBN SGP-E2. Significance was analyzed using the Waller-Duncan test (*P* < 0.05, n = 3).

**Table 1.** Main chemical compositions of SGP-E1.

| SGP-E1 | Protein | Sialic acid | Total sugar |
| --- | --- | --- | --- |
| Content (%) | $65.20 \pm 0.085$ | $11.57 \pm 0.34$ | $16.57 \pm 0.01$ |

### 3.2 N-glycan profile of EBN glycopeptides

The primary components of EBN are Muc5AC mucins, which are rich in sialylated O-glycan and N-glycan chains. The N-glycan profiles of the EBN glycoproteins were determined via DSA-FACE technology, and the electrophoresis peaks were calibrated against the N-glycan standard of serum glycoproteins, as shown in Fig. 3. The N-glycan profile of EBN glycoproteins revealed the presence of ten distinct glycan peaks, the majority of which contained a sialic acid moiety. Specifically, peak 5 is speculated to be A2BG2S1, peak 6 is speculated to be either FA2BG2S1 or FA2G2S1, and peak 7 is speculated to be A2G1. Among the N-glycan peaks of SGP-E1 and E2-HW, FA2BG2S2 exhibited the highest content, whereas A2G1 exhibited the highest content in E2 and E2-LW. Notably, SGP-E1 released N-glycan chains abundant in sialic acid, whereas the majority the N-glycans released from SGP-E2 (peaks 7 and 8) lacked terminal sialic acid.

**Fig. 3.**
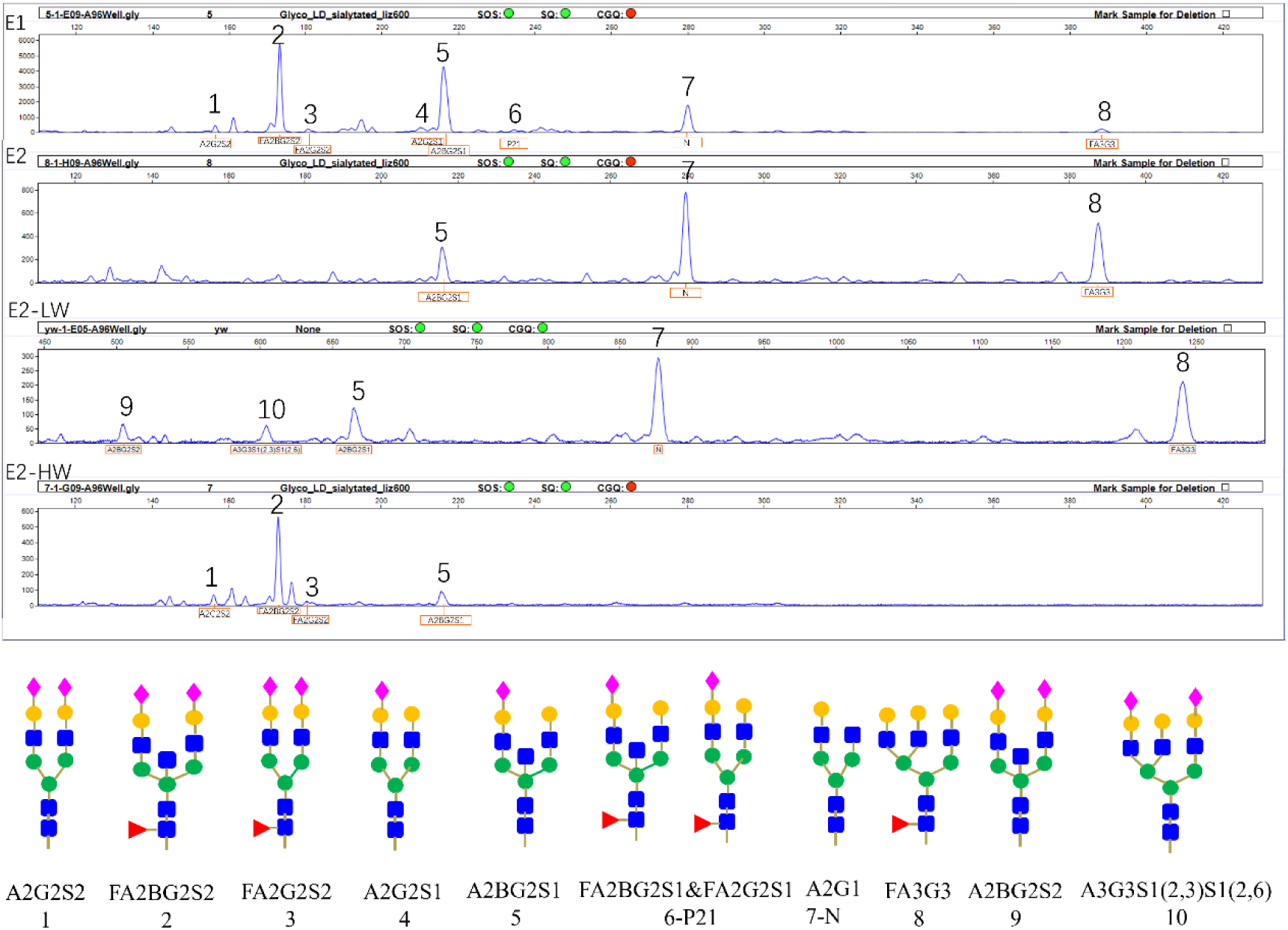
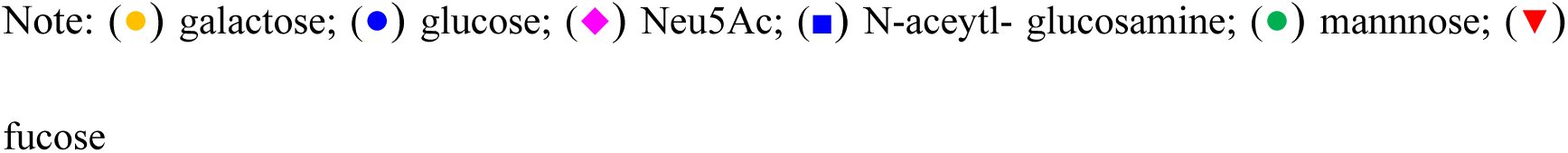
N-glycan profile of EBN hydrolysates obtained by DSA-FACE technique.

### 3.3 Inhibition of C. albicans by dietary sialoglycopeptides

*C. albicans* is a significant opportunistic pathogen commonly found in the human gastrointestinal tract and oral cavity. Infection by this microorganism substantially increases the risk of developing diseases such as oral cancer, atherosclerosis, and diabetes (Aitken-Saavedra et al., 2018; Ayuningtyas et al., 2022; X. Wang et al., 2024). Sialoglycans can bind to surface proteins of *C. albicans*, thereby hindering its attachment to and invasion of host cells (Takagi et al., 2022). In this study, we investigated the effects of EBN glycopeptides and CGMP on the growth of *C. albicans*, with Neu5Ac and 3’-SL as controls (Fig. 4A). The hydrolysates of EBN (SGP-E1, SGP-E2) significantly inhibited the growth of *C. albicans*, particularly the low-molecular-weight portion of E2-LW (*P* < 0.05). When the sialic acid moiety was removed, the inhibitory capacity of Desia-E1 and Desia-E2 was significantly reduced. Interestingly, the high-molecular-weight portion of E2-HW, with a relatively high sialic content, inhibited *C. albicans* to a lesser extent. CGMP and Neu5Ac were unable to fully inhibit the growth of *C. albicans,* whereas 3’-SL had no significant inhibitory effect. Furthermore, EBN SGP-E2 exhibited a dose-dependent inhibitory effect on *C. albicans* (*P* < 0.05), as shown in Fig. 4B. Therefore, the EBN glycopeptide can significantly inhibit the growth of *C. albicans*, with the presence of terminal sialic acid playing a crucial role. However, the inhibitory effects of Neu5Ac, 3’-SL, and CGMP, which also contain sialic acid, on the growth of *C. albicans* were significantly attenuated. Among the EBN hydrolysates, the most pronounced inhibition of *C. albicans* growth was observed in the lower-molecular-weight groups (E2 and E2-LW) compared to the higher-molecular-weight fractions (E1 and E2-HW). Among these groups, E2-LW, despite having a lower sialic acid content than E2-HW, exhibited a stronger inhibitory effect on *C. albicans*. This may be attributed to the fact that the small-molecule E2-LW glycopeptide contains an N-glycosylated chain, which is more likely to stretch out and bind to the receptor on *C. albicans*.

**Fig. 4.**
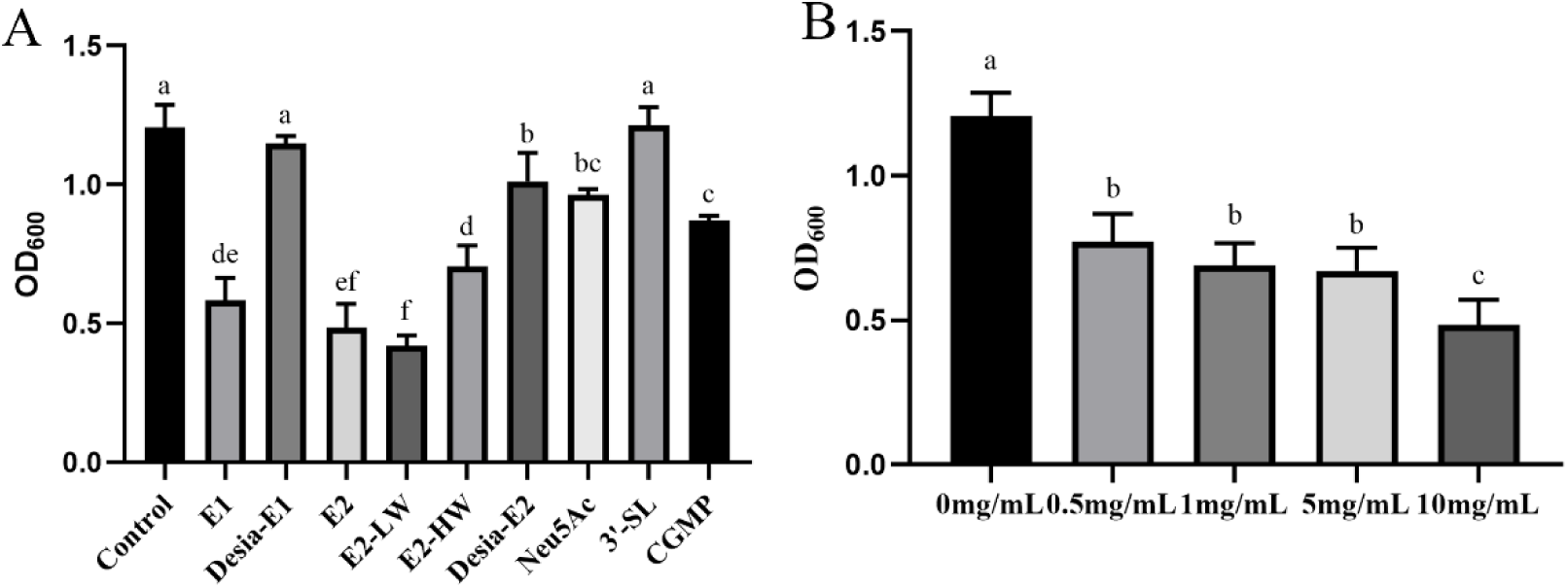
Inhibition of *C. albicans* by various sialoglycans. Note: (A) Different sialoglycans and Neu5Ac (10 mg/L); (B) EBN SGP-E2. Significance was analyzed using the Waller-Duncan test (*P* < 0.05, n = 3).

### 3.4 Inhibition of H. pylori by dietary sialoglycopeptides

*H. pylori* infection is a highly prevalent bacterial infection, with the sialic acid-binding adhesin (SabA) playing a crucial role in its colonization of the oral cavity and gastric mucosa (C. S. Chang et al., 2012; Linden et al., 2008). This study aimed to compare the inhibitory effects of the EBN glycopeptides and CGMP on *H. pylori*, as presented in Fig. 5. Both SGP-E1 and SGP-E2 exhibited similar inhibitory effects on *H. pylori*, demonstrating a dose-dependent inhibition. However, between the two EBN SGPs, SGP-E2 group exhibited a significantly stronger inhibitory effect on *H. pylori* at the same concentration (*P* < 0.05), as depicted in Fig. 5A. Additionally, both SGP-E1 and SGP-E2 exhibited strong inhibitory effects on *H. pylori*, as demonstrated in Fig. 5B. Similarly, E2-LW (molecular weight < 10 kDa) exhibited the strongest inhibitory effect, with an inhibition rate of 37.6%, which was significantly higher than that observed in the other groups (*P* < 0.05). In contrast, the inhibitory capacity of E2-HW was attenuated compared to that of SGP-E2 and E2-LW. The Neu5Ac, 3’-SL, and CGMP glycopeptide exhibited slight inhibitory effects on *H. pylori*, with no significant differences observed among them (*P* > 0.05). Notably, the removal of sialic acid from the EBN hydrolysate, nearly abolished its inhibitory effect on *H. pylori*. In summary, EBN hydrolysates exhibited a dose-dependent inhibitory effect on *H. pylori*, emphasizing the importance of terminal sialic acid in its inhibition.

**Fig. 5.**
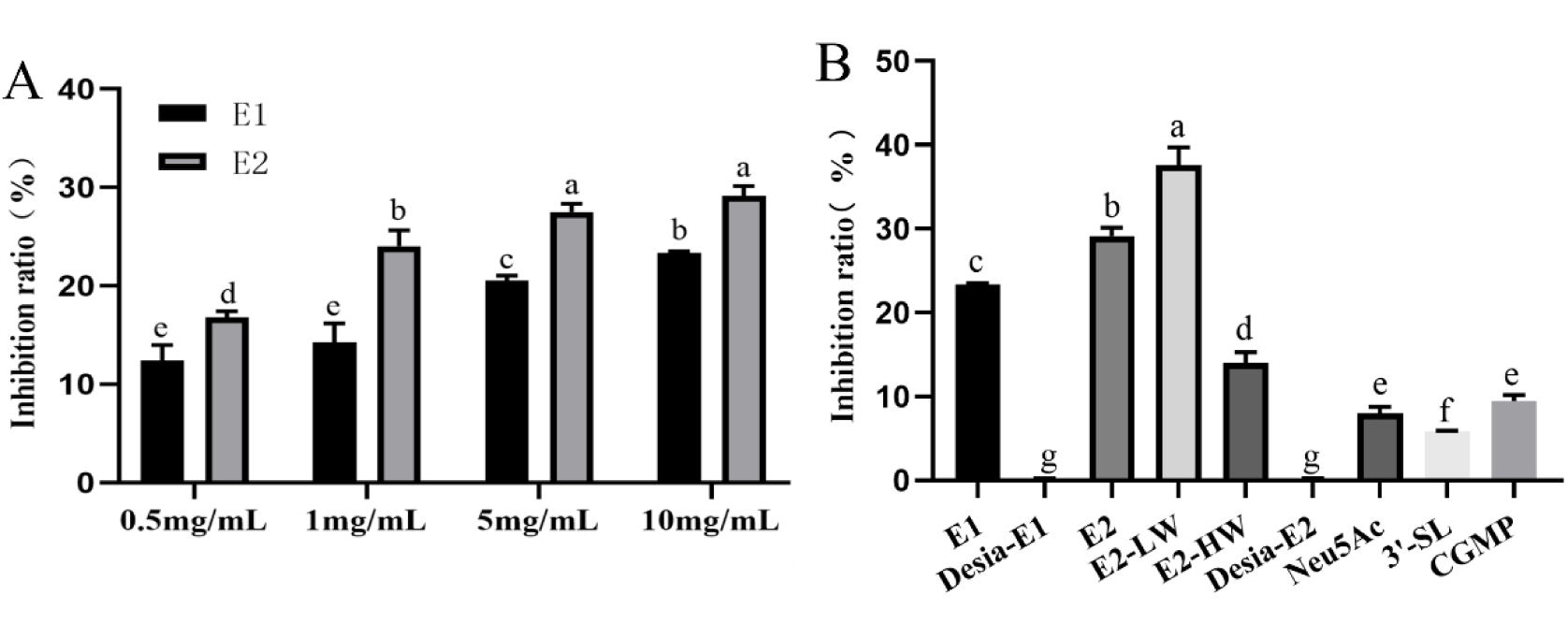
Inhibition of *H. pylori* by various sialoglycans. (A) SGP-E1 and SGP-E2; (B) Different sialoglycans and Neu5Ac (10 mg/mL). Significance was analyzed using the Waller‒Duncan test (*P* < 0.05, n = 3).

### 3.5 Inhibition of avian influenza virus by dietary sialoglycopeptides

Sialic acid serves as a receptor for AIV infection in host organisms and plays a crucial role in host cell invasion (Spackman, 2014). Previous studies have shown that dietary sialoglycans, such as EBN and ovomucin, have the ability to inhibit viral infections (Haghani et al., 2016; Q. Xu et al., 2018). In this study, we employed MAL-II as an AIV mimic to investigate the inhibitory effects of EBN hydrolysates and CGMP, as presented in Fig. 6. Our findings show that MAL-II binding to MDCK cells was significantly inhibited upon the addition of 0.5 mg/mL SGP-E1, compared to the control group, where MAL-II was able to bind to MDCK cells. Similarly, the binding of MAL-II to MDCK cells was also inhibited by 0.5 mg/mL SGP-E2, although the inhibitory effect was less pronounced than that of SGP-E1. Furthermore, our results demonstrate that the inhibitory effect of EBN hydrolysates on the binding of MAL-II to MDCK cells was enhanced with increasing doses of SGP-E1 and SGP-E2, as shown in Fig. 6D and 6E. Notably, when the terminal sialic acid on EBN SGP was removed, the inhibitory effect on MAL-II binding to MDCK cells was significantly diminished (Fig. 6F). Specifically, SGP-E1 most significantly inhibited (*P* < 0.05) the binding of MAL-II to MDCK cells at a concentration of 10 mg/mL, with an inhibition efficiency of 86.28%. In contrast, Desia-E1 and Desia-E2 showed inhibition rates of only 45.83% and 25.62%, respectively. The inhibition efficiency of CGMP derived from milk was 68.95%, which was comparable to that of E2. Moreover, the inhibition rate of the Neu5Ac was only 53.31%, whereas that of 3’-SL was only 26.92%. These findings suggest that the EBN-derived hydrolysate SGP-E1 exhibits the most potent inhibitory effect on MAL-II binding to MDCK cells, highlighting the crucial role of the sialylation structure. The superior anti-AIV efficacy of SGP-E1 may be attributed to its higher abundance of N-glycan chains with sialic acid. For SGP-E2, which exhibited an enhanced antibacterial effect, further hydrolysis with a third protease may fully expose the O-glycan chain.

**Fig. 6.**
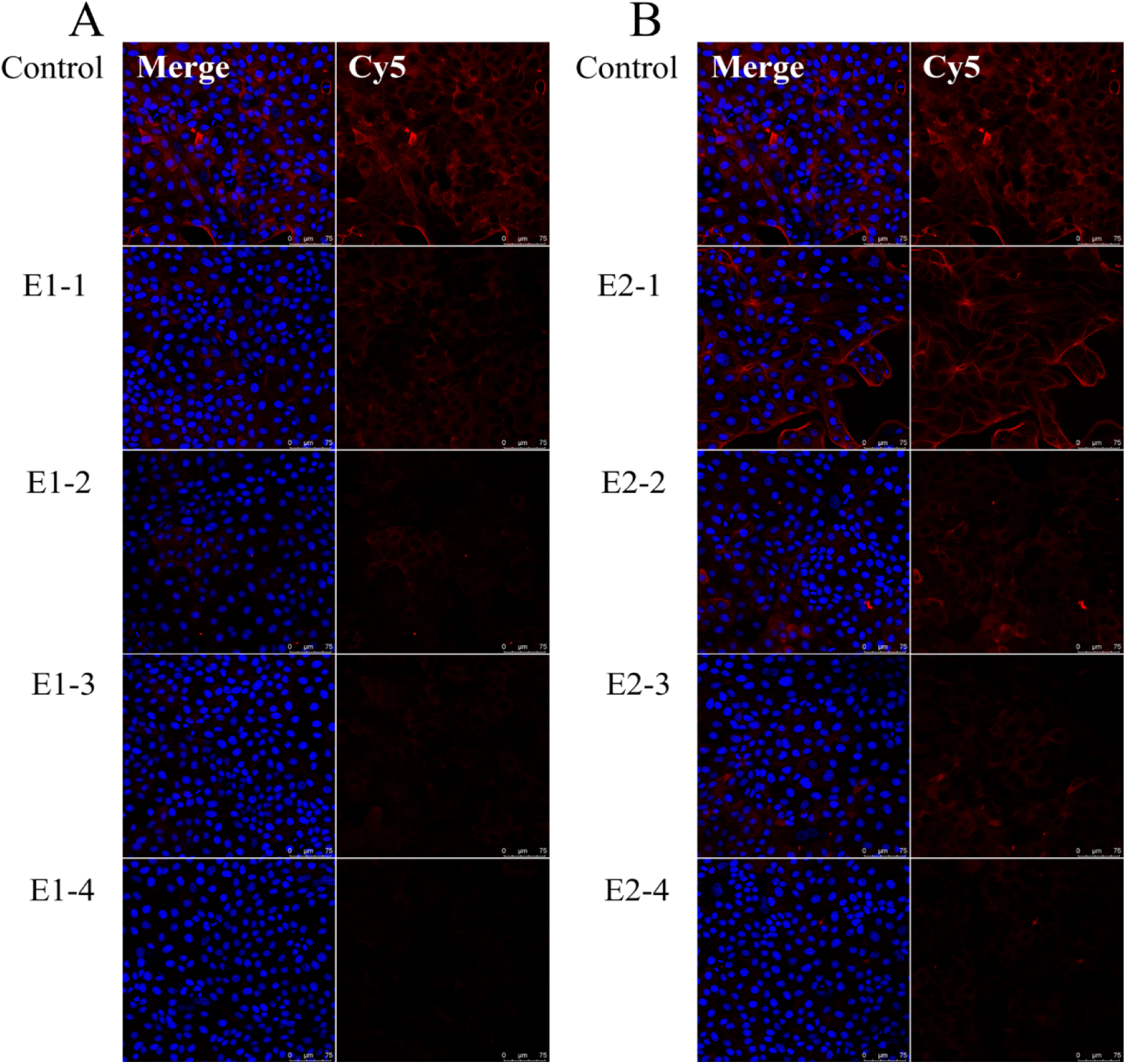
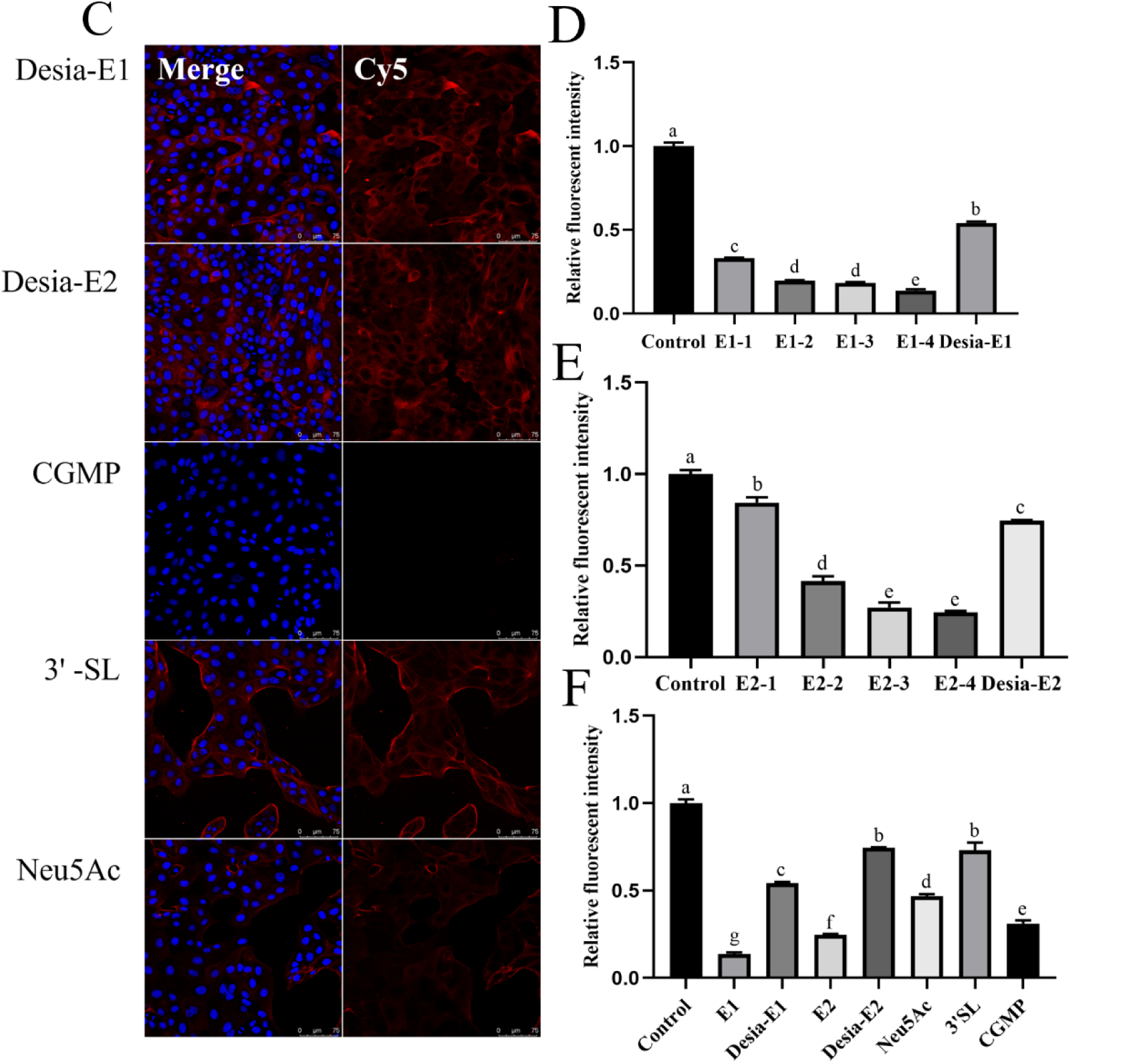
Inhibition of MAL-Ⅱ binding to MDCK cells by various sialoglycans. Note: (A), (B) and (C) Binding of Cy5-labeled MAL-II to MDCK cells in the presence of SGP-E1, SGP-E2, Desia-E1, Desia-E2, Neu5Ac, 3’-SL and CGMP; (D) Relative fluorescence intensity of Cy5-MAL-II binding to MDCK cells in the presence of SGP-E1 (E1-1, E1-2, E1-3, and E1-4 were 0.5 mg/mL, 1 mg/mL, 5 mg/mL, and 10 mg/mL, respectively) and Desia-E1; (E) Relative fluorescence intensity of Cy5-MAL-II binding to MDCK cells in the presence of SGP-E2 (E2-1, E2-2, E2-3, and E2-4 were 0.5 mg/mL, 1 mg/mL, 5 mg/mL, and 10 mg/mL, respectively) and Desia-E2; (F) Relative fluorescence intensity of Cy5-MAL-II binding to MDCK cells in the presence of different sialoglycans and Neu5Ac. Significance was analyzed using the Waller‒Duncan test (*P* < 0.05, n = 3).

### 3.6 Modulation of gut microbiota in the elderly individuals by dietary sialoglycopeptides

#### 3.6.1 Changes in organic acid during in vitro simulated fermentation

Short-chain fatty acids (SCFAs), such as acetic acid, propionic acid, and butyric acid, play an important role in maintaining human health, and the SCFA content in feces is closely correlated with colonic SCFA levels and the composition of the intestinal microbiota (Yamamura et al., 2020). Dietary mucins, such as EBN, can influence the production of organic acids by intestinal flora. The organic acid yields from different groups in vitro anaerobic fermentation with fecal bacteria from elderly individuals are shown in Fig. 7. Acetic acid showcased the highest yield. The yields of lactic acid (Fig. 7A) and SCFAs in glycopeptide E1 group were higher than those in the blank and other control groups (*P < 0.05*). When the terminal sialic acid in glycopeptide E1 was removed, the organic accumulation showed no significant difference between Desia-E1 group and the blank group. This suggests that the bound sialic acid on glycoproteins plays an important role in promoting intestinal microbial fermentation and lactic acid accumulation. Notably, the CGMP group (approximately 7% sialic acid, O-glycosylation) exhibited significantly lower lactate production compared to the E1 group (*P < 0.05*). Additionally, the small molecule 3’-SL and Neu5Ac did not enhance lactate yield. As shown in Fig. 7B and 7C, both acetic acid and butyric acid yields in the E1 group were significantly higher than those in the blank and the Desia-E1 groups (*P < 0.05*). Acetic acid production was significantly higher in the CGMP group than in the blank group. Whereas butyric acid production did not show a significant difference from the blank group, suggesting that CGMP also promotes the production of the SCFA, but not as effectively as the E1 group. The isovaleric acid yield was lower yield in this in vitro fermentation, but the E1 group still exhibited an increase compared to the other groups (Fig. 7D). The results suggest that glycans with distinct sialylation patterns exert differential effects on the production of organic acids by the intestinal microbiota in elderly individuals. Compared to the O-glycosylated CGMP and small molecule 3’-SL and the Neu5Ac, EBN glycopeptide E1 with intact sialyl residues linked to an N-glycan structure was more conducive to the accumulation of organic acids.

**Fig. 7.**
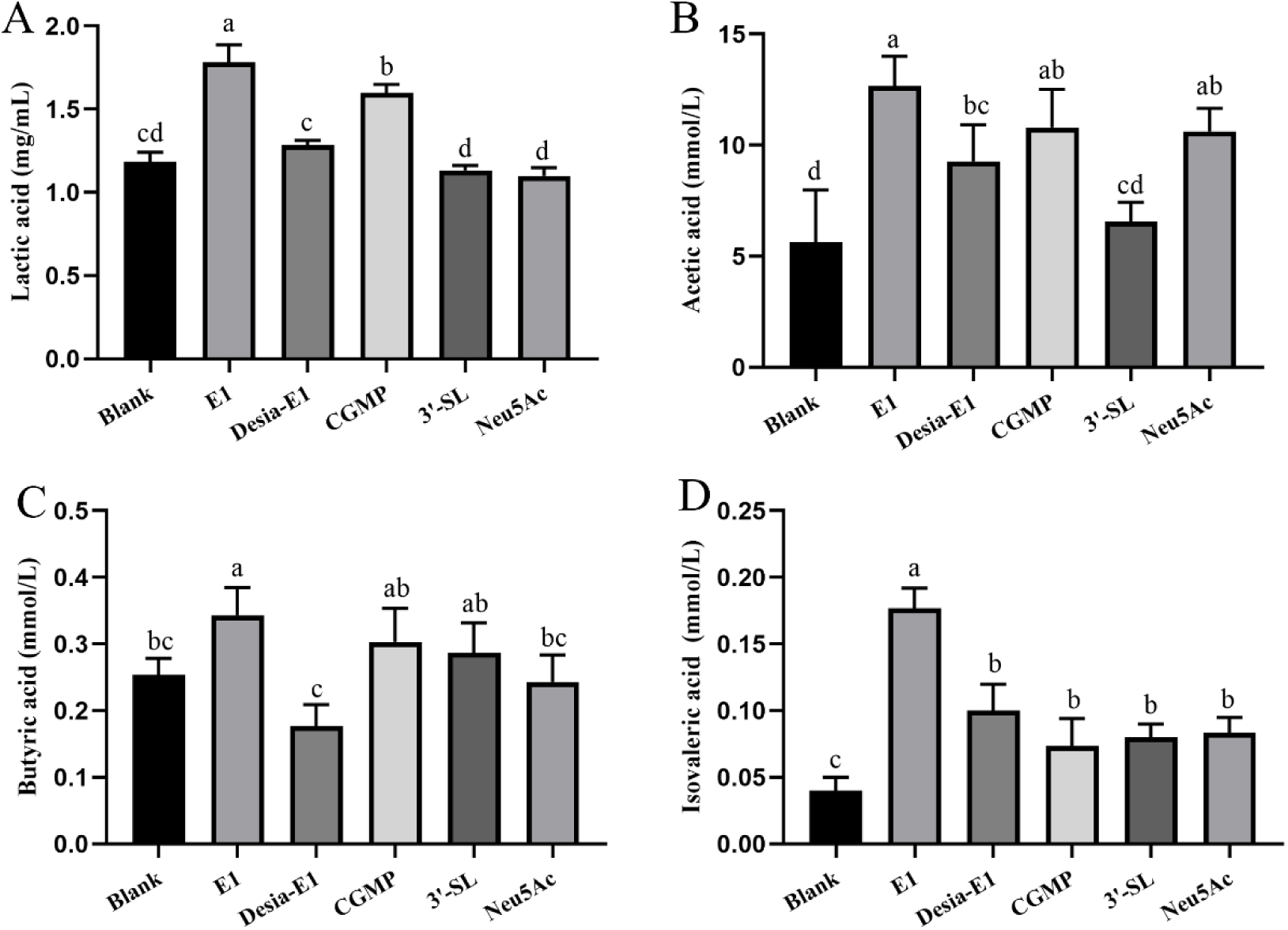
Organic acid accumulation under in vitro fermentation with elderly fecal bacteria (24 h). Note: (A) lactic acid; (B) acetic acid; (C) butyric acid; (D) isovaleric acid.

#### 3.6.2 Abundance Analysis of gut microbiota

Gut microbiota typically refers to the diverse microbial communities within the human gastrointestinal tract that influence both human health and the development of diseases. The distribution and availability of sialic acid and its derivatives within the human gastrointestinal tract can modulate the susceptibility of both gastrointestinal microorganisms and their hosts to inflammation and infection, thereby influencing the composition and abundance of gastrointestinal bacteria as well as pathogen colonization (Bell et al., 2023). We compared the regulatory effects of two glycopeptides on the gut microbiota of the elderly individuals through in vitro fermentation combined with 16S rRNA high-throughput sequencing, and the results are shown in Fig. 8. The top 10 species at the phylum level for abundance analysis was depicted in Fig. 8A, and the top 30 species at the genus level for abundance analysis via heatmap visualization are presented in Fig. 8B.

**Fig. 8.**
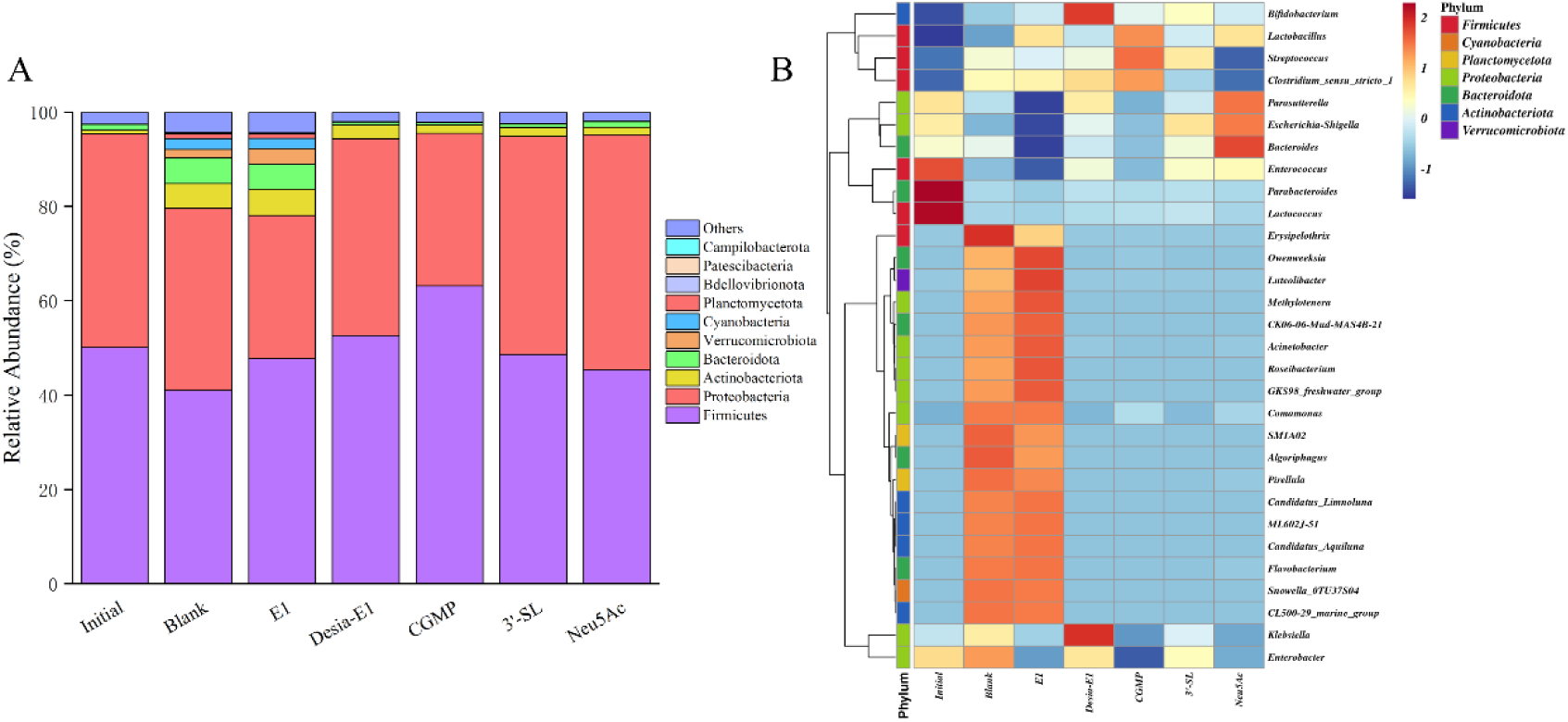
Composition of intestinal microbiota after in vitro fermentation (24 h). Note: (A) Stacked plot of relative abundance at the phylum level (top10); (B) Heat map of relative abundance at the genus level (top30).

At the phylum level, all groups showed an increased relative abundance of Firmicutes compared to the blank group, while the E1 and CGMP groups decreased the relative abundance of Proteobacteria. In contrast, the Desia-E1, 3’-SL and Neu5Ac groups showed the opposite trend. Additionally, the E1 group increased the abundance of Actinobacteria, a highly diverse group of microorganisms known for producing numerous biologically active natural products and clinical drugs (van Bergeijk et al., 2020).

At the genus level, the E1 group significantly increases the relative abundance of *Methylotenera*, *Roseibacterium*, and *Lactobacillus*, while significantly reducing the relative abundance of the harmful bacteria *Escherichia-Shigella* and *Klebsiella*, compared to the blank group. *Methylotenera and Roseibacterium*, in particular, are potential probiotics known for their protective effects on gut health (Hou et al., 2020; Yun et al., 2014). *Escherichia-Shigella* and *Klebsiella* are representative harmful bacteria known for their pro-inflammatory and pathogenic properties (Baltazar-Díaz et al., 2022). In addition, *Lactobacillus* were also enriched in the E1 and CGMP groups compared to the blank group. The Desia-E1 group similarly enriched the common probiotic *Bifidobacterium*, while the abundance of harmful bacterium *Klebsiella* was also increased. The 3’-SL group also increased the relative abundance of *Bifidobacterium*. However, Neu5Ac group increased the abundance of harmful bacteria such as *Escherichia-Shigella*. These results suggest that higher levels of Neu5Ac monomer may lead to intestinal microecological dysregulation, thereby exacerbating the risk of inflammation and infection (Liang et al., 2023). While the bound Neu5Ac showed excellent probiotic properties, the sialylated N-glycans and O-glycans in E1 and CGMP may be more effective in enhancing the abundance of probiotics and reducing the abundance of harmful bacteria. Similarly, N-glycans in milk have been reported to exhibit strong antibacterial and bacteriostatic activity (Yin et al., 2022). In this study, EBN glycopeptide E1 with an N-glycan structure also showed effective inhibition of harmful pathogens.

#### 3.6.3 Analysis of the alpha diversity

Alpha-diversity, also referred to as single-sample complexity, reflects significant variations in the richness and diversity of microbial communities. In this study, the ACE (Abundance-based Coverage Estimator) index, which reflects microbial species richness, and the Shannon index, which reflects microbial diversity, were chosen to investigate the effects of different glycopeptides on the intestinal flora of elderly individuals, as illustrated in Fig. 9. The species richness and diversity of the intestinal flora in the E1 group were significantly increased (*P < 0.05*) compared to the initial flora. However, this index of Desia-E1 group was similar to that of initial flora in terms of both richness and diversity. The CGMP and 3’-SL group did not promote the diversity of the intestinal flora in elderly individuals. In contrast, the diversity of the intestinal flora in the Neu5Ac group decreased compared to the initial group. This suggests that elevated Neu5Ac levels may lead to intestinal microecological dysbiosis(Cotillard et al., 2013). It appears that bound Neu5Ac within the N-glycan structure may have a greater capacity to promote the richness and diversity of the gut flora in the elderly.

**Fig. 9.**
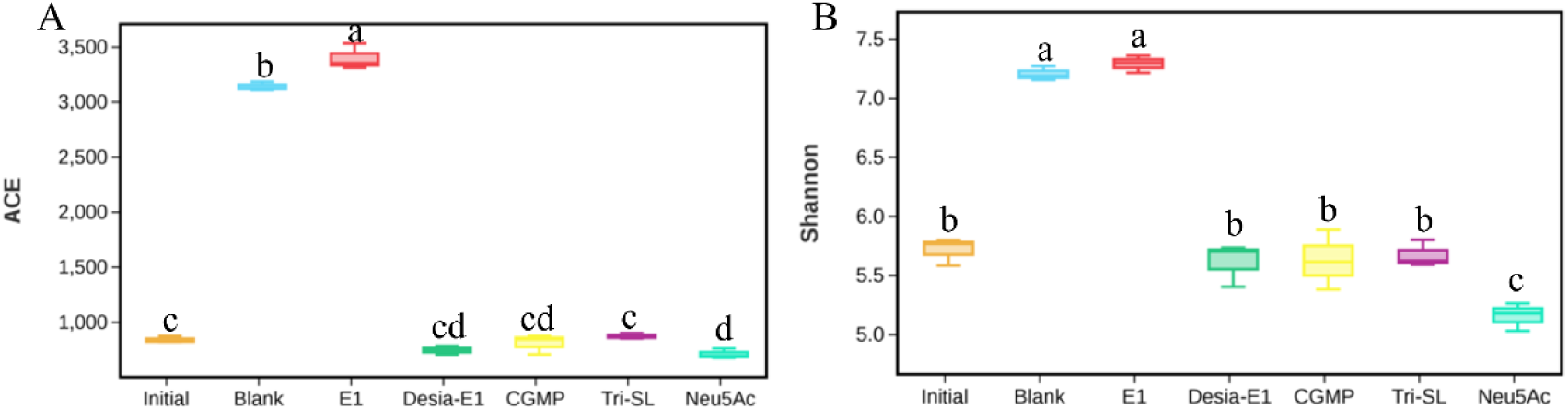
Alpha diversity analysis of intestinal microbiota in different groups based on a 97 % threshold under in vitro fermentation conditions. Note: (A)-ACE index; (B)-Shannon index.

#### 3.6.4 LEfSe analysis

The Linear Discriminant Analysis Effect Size (LEfSe) is used to identify species with significant differences in abundance between populations and can be employed to compare species across different groups. Fig. 10A showcases the species flora that significantly differed between groups (LDA score > 4). Fig. 10B presents an evolutionary branching diagram for the differential species, with taxonomic levels from phylum to genus represented by circles radiating inwardly and outwardly. The relative abundance of the differential species is proportional to the diameter of each circle. Group E1 exhibited a higher abundance of Actinobacteria, which play a key role in maintaining intestinal homeostasis (Binda et al., 2018). Both the probiotic *Lactobacillus* and the harmful bacterium *Clostridia* were enriched in the CGMP group. The harmful bacterium *Escherichia-Shigella* is abundant in the Neu5Ac group. The E1 and CGMP groups show enhanced probiotic effects; notably EBN glycopeptide E1 exhibits remarkable probiotic activity on the intestinal flora of the elderly.

**Fig. 10.**
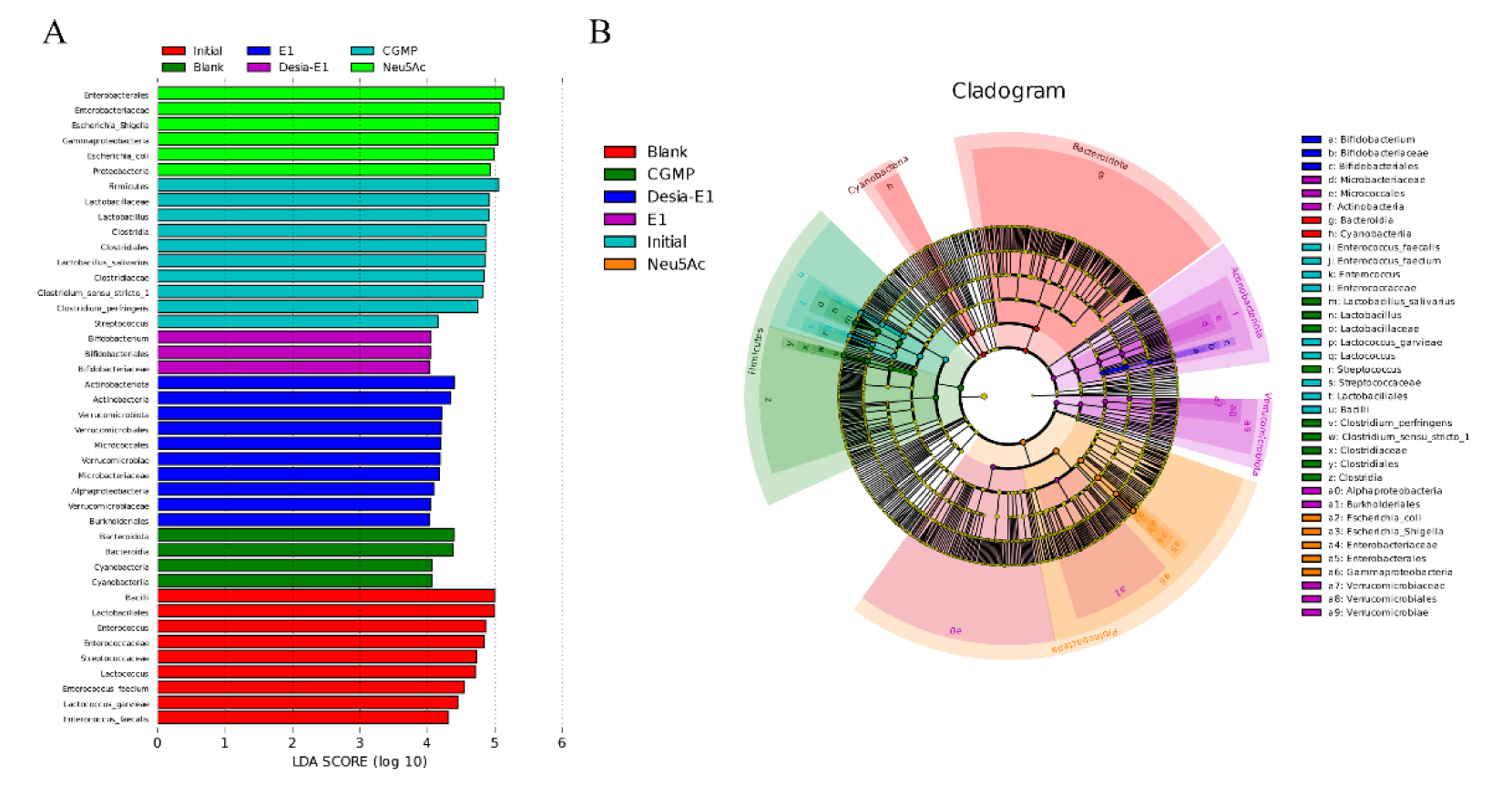
LEfSe analysis of intestinal microbiota in different groups under in vitro fermentation conditions. Note: (A) LDA value distribution histogram; (B) Evolutionary branching diagram.

## 4. Discussion

AIV contains key glycoprotein antigens, namely, hemagglutinin and neuraminidase. These antigens play crucial roles in the virus invasion process by cleaving sialic acid on the respiratory tract of the host (Kumlin et al., 2008; McAuley et al., 2019). Early studies have indicated that AIV utilizes sialic acid linked to galactose via an α2,3-glycosidic bond as a cellular receptor (Olofsson et al., 2005). The expression of α2,3-linked sialic acid on glycoproteins is significantly reduced during pregnancy, thereby increasing the susceptibility of pregnant women to AIV infection (Ding et al., 2021). However, high expression of α2,3-linked sialic acid in the glycoproteins of lactating women may help protect them against AIV infection (Ding et al., 2022). Our study revealed that the hydrolysate of EBN exhibits a stronger inhibitory effect on the influenza virus, with the higher-molecular-weight SGP-E1 (sialoglycopeptide-E1) yielding better results than SGP-E2. However, the inhibitory activity against the virus is significantly reduced when sialic acid is removed. Based of the N-glycan profile, we hypothesized that SGP-E1, which contains more sialylated N-glycans, is more effective in inhibiting the influenza virus. In our research, the inhibitory effect of SGP-E2 on AIV was similar to that of CGMP glycopeptide, which undergoes five O-glycosylations, with approximately 1 to 3 glycans bearing sialic acid termini(Y. Lu et al., 2022). For SGP-E2, treatment of EBN with three proteases resulted in a higher proportion of EBN SGP with a smaller molecular weight, leading to increased exposure of sialylated O-glycans, which may mask the sialylated N-glycans. Additionally, a previous study revealed that 3’-SL (3’-sialyllactose) exhibited effective antiviral activity against 13 AIVs (Pandey et al., 2018). However, our study showed that 3’-SL had an inhibitory effect on AIV simulation, although it was the weakest, even inferior to Neu5Ac (N-acetylneuraminic acid). Our findings suggest that sialoglycopeptides, particularly those rich in exposed and sialylated N-glycans, have greater potential in inhibiting influenza viruses. However, the anti-influenza activity of EBN hydrolysate has not been compared with that of established anti-influenza therapeutics, such as Tamiflu and Zanamivir. Therefore, further in-depth research is necessary in the future.

Sialoglycans can bind to specific receptors on the surface of pathogenic bacteria, leading to inhibitory effects. For instance, the adhesion of *C. albicans* to hosts occurs through the binding of receptors on the bacterial surface to glycoproteins on the host cell surface. The binding sites include mannan or mannose-protein complexes, chitin, and adhesins(Y. G. Kim et al., 2022). One glycopeptide, CGMP, possesses various O-glycan modifications that result in various biological activities, such as the inhibition of bacterial and viral adhesion (Fayed, 2012; Y. Lu et al., 2022). Takagi et al. reported that the O-glycans of mucins can interfere with the invasion process of *C. albicans* into host cells (Takagi et al., 2022). Our study revealed that the EBN hydrolysate exhibited remarkable inhibitory activity against *C. albicans* and *H. pylori*. Among these, the EBN glycopeptide E2-LW, with a molecular weight of less than 10 kDa, exhibited the strongest inhibitory effect, despite its lower sialic acid content. The low molecular weight of EBN SGP may expose more O-glycans, facilitating their extension and enabling them to bind more effectively to receptors on pathogen surfaces. The surface receptor SabA of *H. pylori* binds to host Neu5Ac for adhesion (Salcedo et al., 2013), and some sialylated oligosaccharides, such as 3’-SL, facilitate the binding of *H. pylori* to epithelial cells(Im et al., 2021). However, this study demonstrated that 3’-SL did not exhibit strong inhibitory effects against these two pathogens. We speculate that the maximum concentration of 3’-SL at 10 mg/mL may be insufficient to inhibit both pathogens.

Intestinal flora constitutes a fundamental component of the intestinal microecology, participating in important processes such as the synthesis of essential nutrients and carbohydrate metabolism, while also serving as a key intermediary between diet and host health(Zhou et al., 2020). Common prebiotics, including Galacto-oligosaccharide (GOS), inulin, and Fructo-oligosaccharide (FOS), are utilized to modulate the gut microbiota (Y. Li et al., 2024). EBN comprises approximately 25% carbohydrates, the majority of which are sialylated O-glycans and N-glycans, which are metabolized by gut microbes to yield important metabolites, such as short-chain fatty acids and neurotransmitters, thereby improving host health(Jana et al., 2021). For instance, EBN has been shown to promote bacterial species richness and diversity in the intestinal flora, increase beneficial microbial populations, regulate intestinal dysfunction and bolster immunity in mice(C. Li et al., 2024). In prior studies, we confirmed that hen egg ovomucin can enhance the diversity of the intestinal flora in the elderly(Fang et al., 2024). Similarly, we observed that EBN can also increase the richness and diversity of the intestinal flora in the elderly, increase the abundance of beneficial bacteria and reduce the prevalence of harmful bacteria. Among them, both ovomucin and EBN possess N-glycan structure.

Sialic acid, an important glycan, plays a critical biological role in vertebrates, with its structural pattern significantly influencing its functional properties. Table 2 summarizes the structural characteristics of several sialoglycans used in this study. Both the Neu5Ac monomer and CGMP exhibit inhibitory effects on *C. albicans*, *H. pylori,* and AIV. However these effects are not statistically significant. Several reports have confirmed that 3’-SL exhibits effective inhibitory activity against pathogenic bacteria(Im et al., 2021; J. C. Yang et al., 2013), but the dosage administered in this study did not produce a comparable effect. A high content of sialic acid does not necessarily correlate with enhanced inhibitory effect against pathogenic bacteria or viruses. In this study, we observed that the EBN sialoglycopeptide exhibits excellent inhibitory effects on pathogenic bacteria and AIV, potentially due to its abundant sialylation of both O-glycans and N-glycans.

**Table 2.**
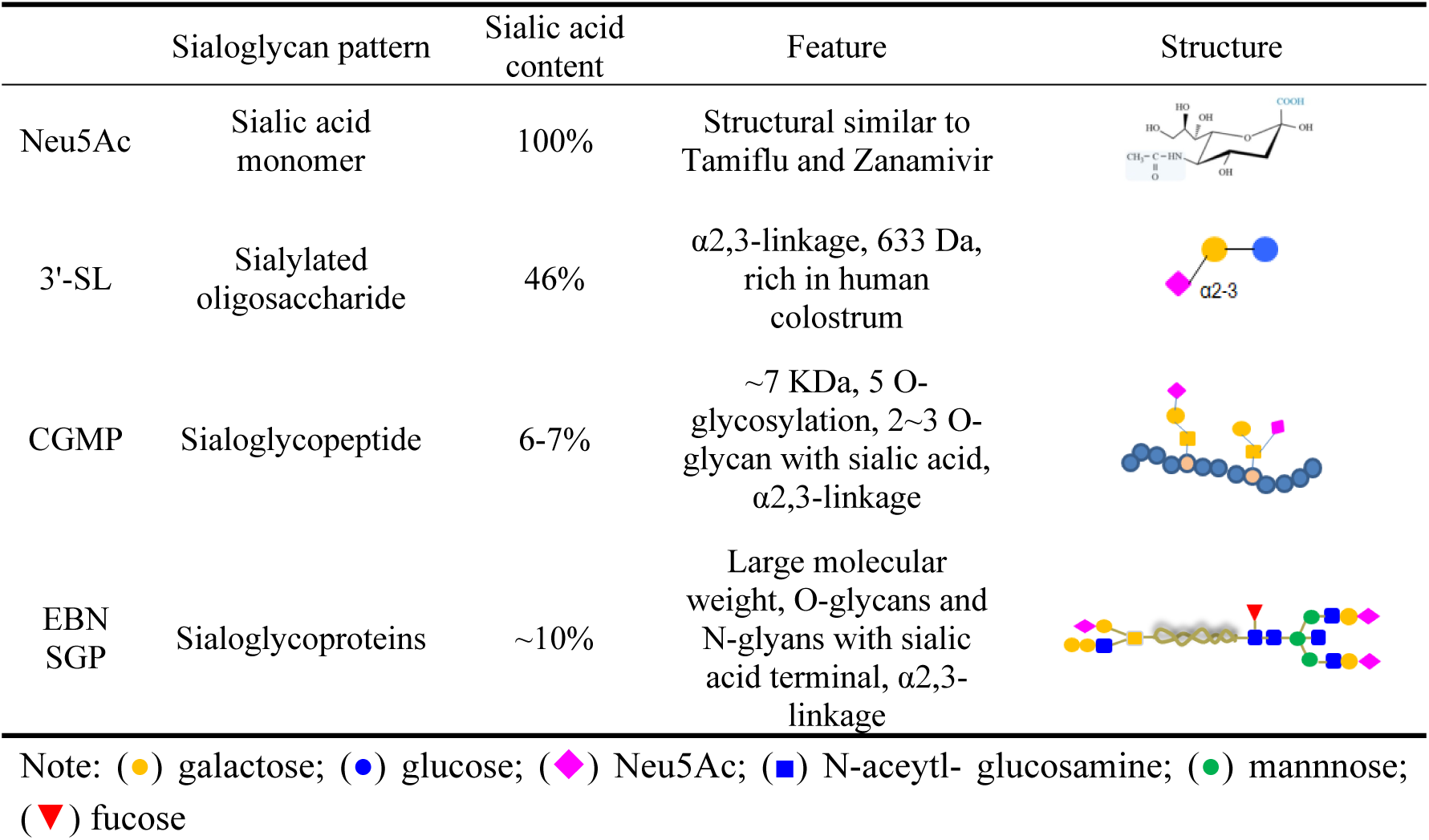
Sialylation patterns of natural sialoglycans.

|  | Sialoglycan pattern | Sialic acid content | Feature | Structure |
| --- | --- | --- | --- | --- |
| Neu5Ac | Sialic acid monomer | 100% | Structural similar to Tamiflu and Zanamivir |  |
| 3'-SL | Sialylated oligosaccharide | 46% | α2,3-linkage, 633 Da, rich in human colostrum | |
| CGMP | Sialoglycopeptide | 6-7% | ~7 KDa, 5 O-glycosylation, 2~3 O-glycan with sialic acid, α2,3-linkage | |
| EBN SGP | Sialoglycoproteins | ~10% | Large molecular weight, O-glycans and N-glycans with sialic acid terminal, α2,3-linkage | |

Note: (●) galactose; (●) glucose; (◆) Neu5Ac; (■) N-acetyl- glucosamine; (●) mannose; (▼) fucose

Currently, reports on the analysis of the N-glycosylation structure of EBN are limited. Early research conducted by Yagi et al. have demonstrated that EBN predominantly contains N-glycan chains with 3 and 4 antennary structures (Yagi et al., 2008). In our study, we utilized DSA-FACE technology to analyze the N-glycan profiles of EBN glycoproteins. Through this analysis, we identified 10 different glycan peaks, several of which contained branched GlcNAc moieties on the core pentose. Among these glycan structures, 8 peaks were found to possess sialic acid termini. An in-depth investigation into the N-glycosylation of EBN from different sources is necessary for future work.

N-glycosylation influences the structure and function of many proteins and provides recognition sites for protein folding and transport, host defense against pathogen invasion and inflammation(Gu & Isaji, 2024). Dietary glycoproteins, with N-glycans exhibiting various structural modifications, such as sialylation, and fucosylation, can elicit distinct biological functions (Zhang et al., 2024). Previous studies have shown that removal of fucose in the N-glycans of bovine milk proteins significantly reduces their antibacterial activity (W. L. Wang et al., 2017). Sialylation of N-glycans on glycoproteins have been shown to not only prevent influenza virus-induced hemagglutination, but also inhibit the adhesion of pathogenic *Escherichia* to epithelial cells(Hou et al., 2020). Sialylation of N-glycans in cow lactoferrin facilitates the binding and sequestration of lipopolysaccharide (LPS) (Latorre et al., 2010). N-glycans with both sialylation and fucosylation modifications in lactoferrin have been shown to significantly inhibit the growth of *Staphylococcus aureus*(Du et al., 2018). In this study, glycoproteins and glycopeptides derived from EBN had strong inhibitory effects against pathogenic microorganisms, including *C. albicans*, *H. pylori* and AIV. Ten different structures of N-glycans were identified in EBN. Among these, eight N-glycans had sialylation modification, four had fucosylation moiety, and three had both sialylation and fucosylation modifications. In EBN SGP E1, a higher content of sialylated N-glycan was exposed, whereas in EBN SGP E2-LW, more fucosylated N-glycans were exposed. These results indicate that, in EBN glycoproteins, N-glycans with fucosylation modification play a crucial role in inhibiting *C. albicans* and *H. pylori*, while N-glycans with high levels of sialylation, as well as those possessing sialylation and fucosylation modifications, are advantageous for inhibiting AIV.

Sialic acid is essential for the biological functions of EBN glycoproteins. These glycoproteins and glycopeptides, with different molecular weights, show various inhibitory effects on pathogens, likely due to their structural integrity. Therefore, analyzing the fine structures of sialoglycoproteins and sialoglycopeptides with varying molecular weights is crucial for future research. Specially, the biological function of N-glycosylation in dietary glycopeptides and its synergistic interaction with O-glycosylation require further investigation.

## 5. Conclusion

Sialoglycopeptides have been demonstrated to potently inhibit pathogenic bacteria and avian influenza virus (AIV). The sialylation pattern of these glycopeptides plays a vital role in mediating their inhibitory effects. Within a specific concentration range, EBN SGP-E2 glycopeptides with lower molecular weights exhibited stronger inhibitory effects on the growth of *Candida. albicans* and *Helicobacter. pylori*. In comparison to CGMP, Neu5Ac, and 3’-SL, SGP E2-LW exhibited more potent inhibitory effects on both pathogens. Moreover, SGP-E1 and SGP-E2 had dose-dependent inhibitory effects on AIV, with SGP-E1 exhibiting stronger inhibitory effects compared to the CGMP, Neu5Ac, and 3’-SL. This effect may be attributed to the N-glycan structure, which contains a higher sialic acid content in the EBN SGP. Compared to the CGMP, 3’-SL and Neu5Ac groups, SGP-E1 significantly enhanced the production of lactate and SCFAs during in vitro fermentation using fecal bacteria from elderly individuals. SGP-E1 significantly promoted the species richness and diversity of the gut microbiota, enhancing the relative abundance of beneficial bacteria, such as *Lactobacillus* and *Roseibacterium*. Additionally, it reduced the relative abundance of harmful bacteria, including *Escherichia-Shigella* and *Klebsiella*. N-glycan profiling identified 10 distinct glycan peaks, the majority of which contained sialic acid. Compared to the O-glycosylation of CGMP, the N-glycosylation of EBN, which includes sialic acid and fucose moieties, can more effectively inhibit pathogenic microorganisms and enhance their regulatory effects on the intestinal microbiota of elderly individuals. These findings advance our understanding of the relationship between glycosylation structure and the biological functions of sialoglycopeptides.

## CRediT author statement

**Jiao Zhu**: Investigation, Methodology, Writing original draft; **Tiantian Zhang**: Software, Validation; **Jianrong Wu**: Conceptualization, Validation, Writing-Review and Editing; **Hongtao Zhang**: Supervision, Validation.

## Acknowledgements

This work was supported by the National Key Research and Development Program of China (2023YFA0914303).

